# Traditional practices, knowledge and perceptions of fire use in a West African savanna parkland

**DOI:** 10.1101/2020.09.24.311209

**Authors:** Esther Ekua Amoako, James Gambiza

**Affiliations:** Department of Environmental Science, Rhodes University, P.O. Box 94, Grahamstown, South Africa; Department of Ecotourism and Environmental Management, University for Development Studies, P.O. Box 1882 Tamale, Ghana

**Keywords:** socio-cultural practices, fire use, fire control, fire regime, Guinea savanna

## Abstract

Understanding people’s fire practices, knowledge and perceptions of the use of fire and fire regimes can inform fire management plans that could contribute to sustainable savanna conservation and management. We investigated the frequency of fire use, control and perceptions of fire regime for selected livelihood and socio-cultural activities in six districts in the Guinea savanna of Ghana. The majority of respondents (83%) across the study districts indicated that they used fire once a year for at least one of the following activities: land preparation, weed/grass/pest control, burning stubble after harvest, bush clearing around homesteads, firebreaks, charcoal burning and hunting. The study showed a higher frequency of fire use in the dry season for land preparation for cropping. Less than a fifth of the respondents (17%) indicated that they do not use fire for any of the above activities. The majority of respondents (62%) across the districts mentioned that they controlled their use of fire to prevent destruction to property, with the remaining 3% who indicated the prevention of killing or injuring humans. The study showed a higher frequency of fire use for land preparation for cropping than for the other socio-cultural activities. However, respondents rated season of burning as the most important attribute, with little attention to the other attributes of a fire regime, contrary to what is theoretically recognized. Understanding traditional fire use practices in terms how to regulate the mix of frequency, intensity/severity, season, size and type of fire for these and other socio-cultural purposes could enhance sustainable savanna conservation and management. There is a need to unravel the specifics of fire assisted socio-cultural practices and fire regimes in West Africa.

## 1. INTRODUCTION

Historically, humans have extensively used fire for various land use practices: agriculture, hunting, foraging and pasture management, which has created diverse habitats and landscapes (Sara and Henrik 2002; Luoga, Witkowski and Balkwill 2004; Eriksen 2007; López-Merino *et al.* 2009). The use of fire may be prescribed, controlled, uncontrolled or unintentional, depending on the source of the fire and the purpose of use (Shaffer 2010; Fernandes *et al*. 2013). Burning for agricultural purposes is a common practice worldwide and is mainly used in the tropics (Eastmond and Faust 2006; Tomson, Bunce and Sepp 2015; Hlaing, Kamiyama and Saito 2017).

Thus the practice of fire use in traditional agriculture (crop and animal husbandry) has contributed to the transformation of landscapes over centuries and has been the source of food supply for both rural and urban economies of most countries of sub-Saharan Africa (Halstead 1987; Bassett, 1988; Holden 1993; Bassett *et al*. 2003).

In fire-prone West African savannas, the use of fire is integral in socio-cultural practices that are important for rural livelihoods and socio-economic activities (Rose Innes 1972; Archibald and Bond 2003; Dwomoh and Wimberly 2017). Nonetheless, the savannas provide the land resources where the bulk of cereals, grains and livestock are produced, usually by means of traditional methods of agriculture where fire is used to clear the land of vegetation for cropping, to burn stubble after harvest, as well as for weed management (Amissah, Kyereh and Agyeman 2010; Nyongesa and Vacik 2018). Also, pastoralists in these savannas have often used fire to control pests and to stimulate fresh grass growth in the early part of the dry season (Bassett *et al*. 2003; Coughlan 2013).

Several studies have shown that most rural economies still depend on hunting as a traditional livelihood to supplement household income and protein needs (Adongo *et al*. 2012; Lindsey *et al.* 2013; Day *et al*. 2014; Nyongesa and Vacik 2018). The primary method of hunting in the savanna is the use of fire to flush out animals or by burning early in the dry season to attract animals to be hunted (Walters, Touladjan and Makouka 2014). Most of these fires are difficult to control, especially during the Harmattan (North-Easterly wind) which is characterized by very low humidity and windy conditions so that if fire left uncontrolled, can result in large and persistent fires (Adongo et al. 2012; Abukari and Mwalyosi 2018). However, other studies have reported that fire is also caused by arson, careless disposal of cigarettes and children trying to mimic parents who use fire (Shaffer 2010; Archibald 2016; Nyongesa and Vacik 2018).

The extensive knowledge, values, practices and perceptions of the traditional use of fire in subsistence agriculture, as practised in tropical savanna, particularly in Africa have been underestimated and have received limited attention. Several studies have focused on the management and impact of fire on the environment (e.g. Russell-Smith, Start and Woinarski 2001; Beringer *et al.* 2007; Bilbao, Leal and Méndez 2010; Bento-Gonçalves *et al*. 2012; Seijo, Cespedes and Zavala 2018). However, only a few studies have focused on understanding people’s knowledge, practices and their perceptions of these ‘fire-assisted’ livelihood activities (e.g. Mistry 1998; Eriksen 2007; Shaffer 2010; Fernandes *et al.* 2013; Nyongesa and Vacik 2018).

Understanding how people perceive the role of fire in their socio-cultural and livelihood activities is necessary to inform fire management plans that are well-matched with local knowledge, fire use practices, as well as traditional land use systems (Raish, González-Cabán and Condie 2005; Archibald 2016; Mistry, Bilbao and Berardi 2016). This could contribute to sustainable savanna conservation and management. Thus, the objective of the study was to investigate peoples’ knowledge, perceptions and cultural uses of fire in the Guinea savanna. In this paper, we discuss the frequency of fire use on selected cultural and livelihood activities, fire control practice and the perceptions of fire regimes as defined by Bond and van Wilgen (1996) and Keeley (2009) in high, moderate and low fire frequency districts in the Guinea savanna, Ghana.

## 2 METHODS

### 2.1 Background and setting of the study area

Traditional fire use is perceived as an integral practice in most rural savannas around the world that needs attention. Several studies have shown fire effects on the environment (Dwomoh and Wimberly 2017; Nyongesa and Vacik 2018). Author observation of twenty-five years living in the savanna region of Ghana, shows the widespread use of fire in this ecological zone. In consonance with Eriksen 2007’s research question on ‘why do they burn the bush’ surveys were used to gather information on people’s perception on fire use in rural communities in the north of Ghana in a savanna where farming is the predominant economic activity. Thus, even the few people, who are not directly involved in farming, use fire to for other purposes including clearing bushes around their homes.

The study was conducted in the Guinea savanna ecological zone, Ghana (9.5439° N, 0.9057° W). The climate of the region is tropical with a unimodal rainfall distribution with an annual mean of 1 100 mm (Ghana Meteorological Service 2017). Thus, the region has only one cropping season (Gyasi 1995). The peak of the rainy season ranges between July and September with the rainfall exceeding potential evaporation over a relatively short period (Boubacar *et al.* 2005). The mean annual temperature is 27°C. The region records comparatively higher annual potential open-water evaporation of 2 000 mm than the south, which records 1 350 mm. Due to its proximity to the Sahel, the region experiences dry, dusty northeasterly winds (Harmattan) between November and April, which facilitates the annual vegetation burning (Kranjac-Berisavljevic *et al.* 1999).

The area is characterised by large areas of natural pastures (FAO/UNEP 1981; Raamsdonk *et al.* 2008) with grass species from the sub families of Andropogoneae and Paniceae (Andersson, Kjøller and Struwe 2003; Bocksberger *et al.* 2016), interspersed with fire and drought-resistant woody species from the families of Fabaceae and Combrataceae (Bagamsah 2005; Amoako *et al.* 2018).

#### 2.1.1 Population and social structure

The Northern region of Ghana has a population of nearly 2.5 million, representing ten percent of the total population of the country (Ghana Statistical Service 2010). The population density is relatively low with 37 persons per km^-2^ (Ghana Statistical Service 2010). The majority of the population lives in rural areas. The region has five paramount chiefdoms (traditional areas), namely Mamprusi, Dagomba, Gonja, Nanumba and Mo. Each traditional area represents a major ethnic group in the region (Jönsson 2007). The population of these groups varies, with the largest group being the Dagomba which constitutes 30% of the region’s population. The Mamprusi and Gonja groups comprise over 7% of the population (Ghana Statistical Service 2010).

#### 2.1.2 Agricultural systems

Agriculture contributes to more than 90% of household incomes and employs more than 70% of the population in the region (Ghana Statistical Service 2010). The farming system is traditional agroforestry, which is usually a combination of growing food crops and keeping animals for multiple purposes. Crops are cultivated for about five to ten years among indigenous economic tree species, a system of agriculture which is commonly referred to as agroforestry parklands (Boffa 1995). The crop fields/parklands are allowed to rest for a fallow period of about three to five years to replenish soil fertility after several years of continuous cultivation. Farms are located around one to six km from the compound house (Karbo and Agyare 1999).

Among the major crops grown are maize (*Zea mays*), millet (*Panicum miliaceum*), rice (*Oryza sativa*), yam (*Dioscorea spp*.), and various pulses and vegetables (Gyasi, Kranjac-Berisavljevic, Blay 2004; Brookfield and Gyasi 2009). Cropping is also done around the home compounds in the rainy season and in valleys or along water bodies, especially in the dry season (Gyasi *et al.* 2004). These compound farms are usually permanent, because the soils are replenished by the continuous supply of household waste and manure from livestock (Karbo and Agyare, 1997). They are also burned annually to control ticks and reptiles, as well as for visibility purposes (Gyasi, Kranjac-Berisavljevic and Blay 2004). There are considerable areas of parkland which have not been cultivated, either because of low soil fertility or they have been left fallow for a very long period (Oppong-Anane, 2006).

The Northern region is leading in livestock (ruminants and poultry) production in the country. Livestock is kept on both free range and semi-intensive systems by households and hired herdsman (Fulani), respectively (Karbo and Agyare 1997). Other socio-economic activities are agro-processing (e.g. rice and groundnut processing) and the processing of non-timber forest products (NTFPs) (e.g. shea butter processing).

### 2.2 Data collection

Both stratified and simple random sampling techniques were used. Data on daily fire counts (detected by sensors on Earth observation satellite) from 2013 to 2017 on Ghana, were received from EORIC (Earth Observation Research and Innovation Centre), Ghana in collaboration with the Advanced Fire Information System (CSIR, Meraka Institute), South Africa. Eighteen districts in the Northern region of Ghana, with data on fire counts from 2013 to 2017, were categorised into high fire frequency (6213-15254 counts), moderate fire frequency (2804-6213 counts) and low fire frequency (487-2804 counts) districts (Figure 1).

**Figure 1:**
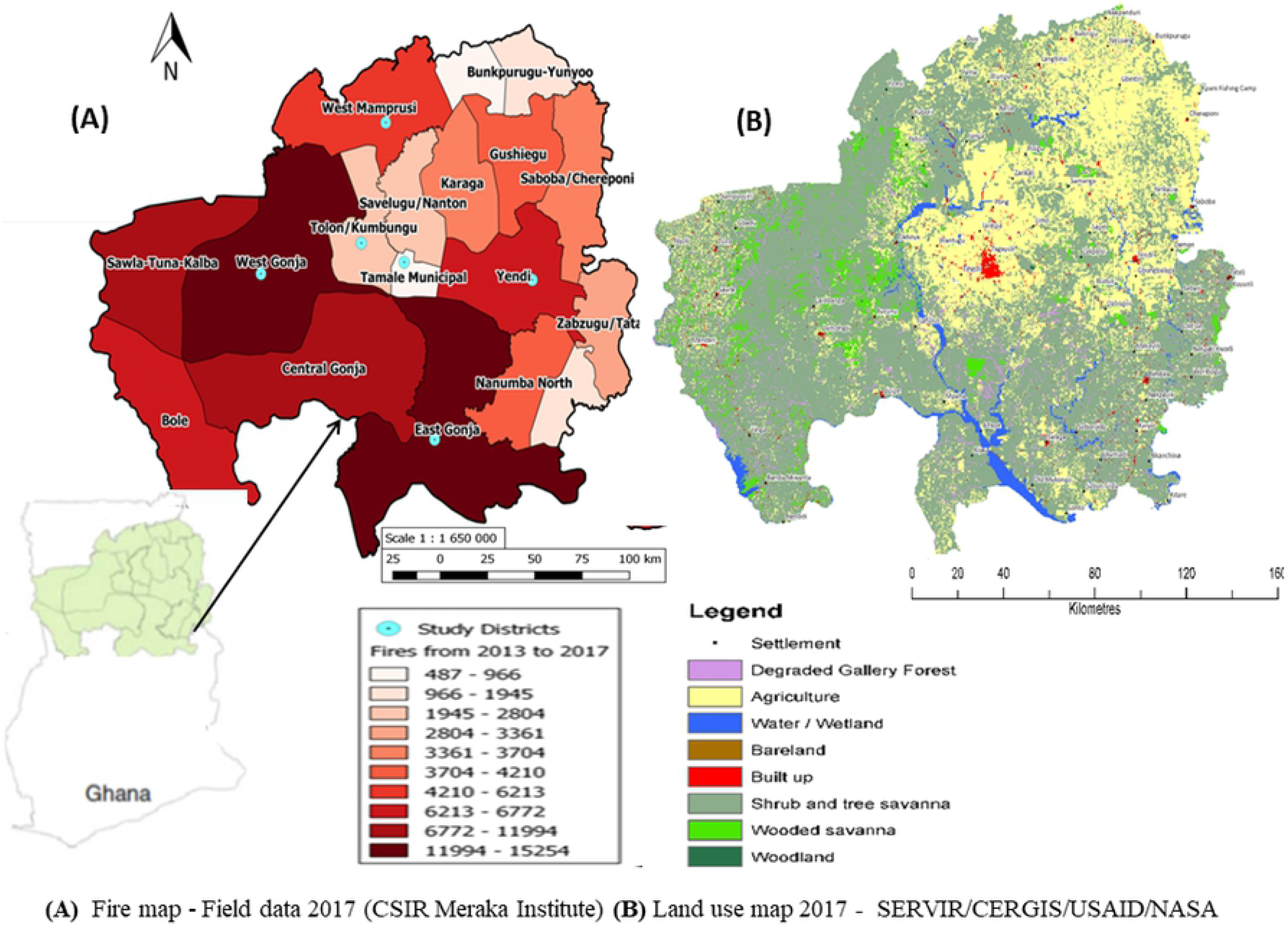
Map of the study area showing the fire frequency gradient (colour gradient), and selected study districts B. Land use map of study area for 2017 (Figure to be printed colour).

Of the 18 districts, six were selected for the survey, in which most of the communities were randomly selected. The communities were Damongo Canteen and Mognori (West Gonja district), Kpalbe and Kushini (East Gonja District) were selected in the high fire frequency districts, whereas Wungu and Kata-Banawa (West Mamprusi district), and Kpligini (Yendi district) are in the moderate fire frequency districts and Jagriguyilli and Nwodua in the Tolon Kumbungu district, Tugu (Tamale South Municipality) were classified as the low fire frequency districts. Tugu and Damongo Canteen are peri-urban communities, while the remaining eight are rural communities (Figure 2)

**Figure 2:**
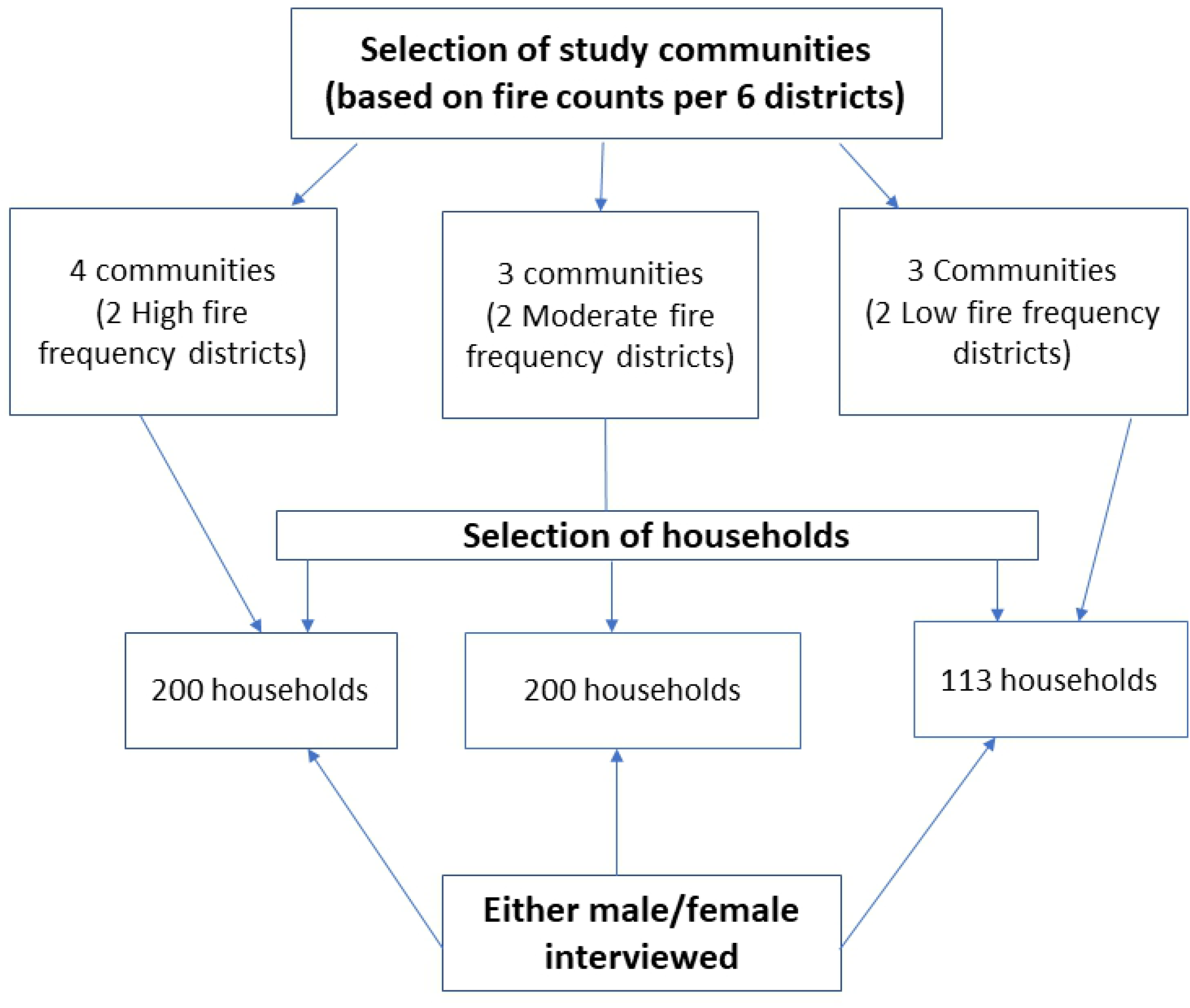
Selection of communities and households for the survey

A survey was conducted by using a questionnaire drafted in English. People were asked questions on their perceptions, practices and knowledge of the use of fire on selected livelihood activities. The questionnaire had three main sections: demographic characteristics, knowledge and perceptions of the use of fire. The questions were mostly closed-ended as opposed to open-ended. The closed-ended questions were mostly three-point (Jacoby and Matell 1971), and five-point Likert type questions as well as binary type questions to measure the importance and frequency of fire use (McDonald 2014). Participants were given options to explain their choice of answers to some of the questions, to aid in guiding responses and facilitating data coding. The question yielded mainly nominal, ordinal and a few continuous variables (age and household size).

The open-ended questions were used to ascertain the activities for which people use fire, people’s practices and knowledge of the use of fire, the impacts of fire on the environment, as well as their knowledge of fire management. The activities (land preparation, hunting, firebreaks, bush clearing around homesteads, and burning stubble after harvest) were selected based on literature on fire use in savannas (Nyongesa and Vacik 2018). The data were collected with the assistance of a research team selected from the University for Development Studies in Tamale, who are fluent speakers of the language spoken in the selected districts. The research team were all males and comprised of five assistants for East and West Gonja Districts, four for West Mamprusi and five for Tolon, Kubungu, Yendi and Tamale South districts. They were trained on the objectives and some terminologies of the study. Households were selected randomly; however, either a male (mostly heads of households) or a female who was willing to answer the questions on fire use was interviewed in a household in each of the communities. Stern (2000) asserts that laws and government regulations are factors that may influence behaviours in responding to questions concerning themselves. Thus, we asked general questions on some sensitive items on hunting and charcoal making and sometimes probing further to get some details. Data were collected from 11 March 2017 to 30 June 2017.

#### 2.2.1 Ethics statement

Formal Ethics Approval was not obtained. The research presents no risks of harm or no effects to the respondents, and involved no formal procedures for which written consents were required. However verbal consents were sought from the community leadership by following community entry protocols via communal meetings and presenting kola nuts to the chiefs, the assembly persons and other leaders in the communities. Some of these communities the research team (particularly the lead author) have worked for several years in different domains of research other than anthropogenic fires. We informed the community members that participation was voluntary, and should anyone feel uncomfortable with any of the questions they could discontinue answering the questionnaire without any negative consequences. We also informed the data will be analysed anonymously and all the information will be stored in a secure space where only the lead researchers and the supervisor have access. Participants were made to understand that the responses were for an academic purpose and I showed my Identity Card to the assembly persons as a proof of my status as a Student. Participants were also assured of confidentiality. All the interviews were conducted in the local dialects of the selected districts as the majority of the respondents could not speak and understand English.

### 2.3 Data analysis

To provide an overview of socio-demographic characteristics, perceptions, knowledge and practices and respondents’s use of fire, data were analysed using both descriptive and inferential statistics. Questions that yielded no and yes answers with reasons and some open-ended answers were classified into themes. All data were coded based on the commonalities in responses and data were entered, cleaned and analysed using SPSS Statistical Software Version 20.0. The total number of responses was 532 of which 295 were from female and 237 were from male respondents. Some of the five-point Likert items were recoded into three-point items to allow more responses in the Likert items for statistical testing. Descriptive statistics were computed and the chi-square test was used to examine the presence of significant relationships among variables across districts (McDonald, 2014). A Pearson’s correlation between people’s perceptions of fire effect on the environment and the frequency of fire use were also computed. Not much data on the open-ended question is presented in this paper therefore, this study should be regarded as an inquiry of how often fire is used for some socio-cultural activities and peoples’ perceptions of a fire regime in the Guinea savanna.

## 3.0 Results

### 3.1 Socio-demographic characteristics of respondents

The highest number of respondents (79%) indicated they were farmers, with only 6% being students; and the remaining 15% engaged in other forms of activities. The mean age of respondents was 39±15 years, with a minimum of 18 years and maximum age of 94 years. About a quarter of the respondents, which were in the majority (21%), were between 26 and 32 years and the lowest (8%) were within the age brackets of 47 and 53 years. The majority of respondents (66%) have no formal education and less than a quarter of the total number of respondents (15%) had education till junior high and senior schools (Table 1).

**Table 1:**
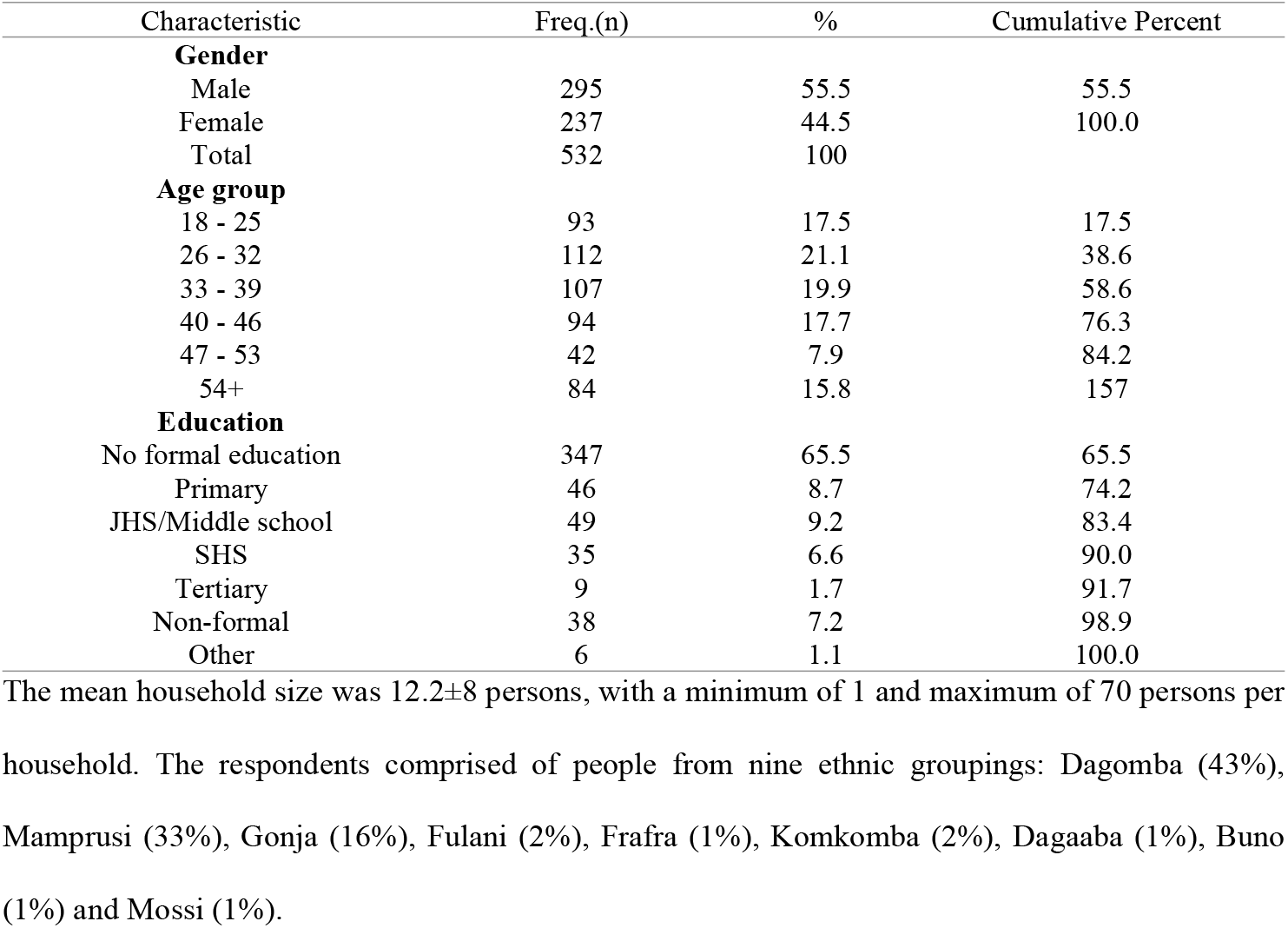
Gender, age and educational distribution of respondents

### 3.2 The frequency of fire use for selected activities

The majority of respondents (83%) across the study districts indicated that they used fire for at least one of the selected activities: land preparation, weed/grass/pest control, burning stubble after harvest, bush clearing around homesteads, firebreaks, charcoal burning and hunting (Table 2). Less than a fifth 17% of said that they do not use fire in these activities. There were varied responses on how often fire was used for the selected activities across the high, moderate and low fire frequency districts. There was a significant association between fire frequency and land preparation for crop production amongst the three categorised districts (χ2 = 39.5, df = 9, p < 0.001). The majority of respondents in the high (86%), moderate (85%) and low (75%) fire frequency districts indicated they used fire once a year for land preparation, while as low as 8%, in both high and moderate and 2% in low fire frequency districts, used fire twice a year for land preparation (Table 2). On average, 14% of the respondents across the districts never used fire for land preparation.

**Table 2:**
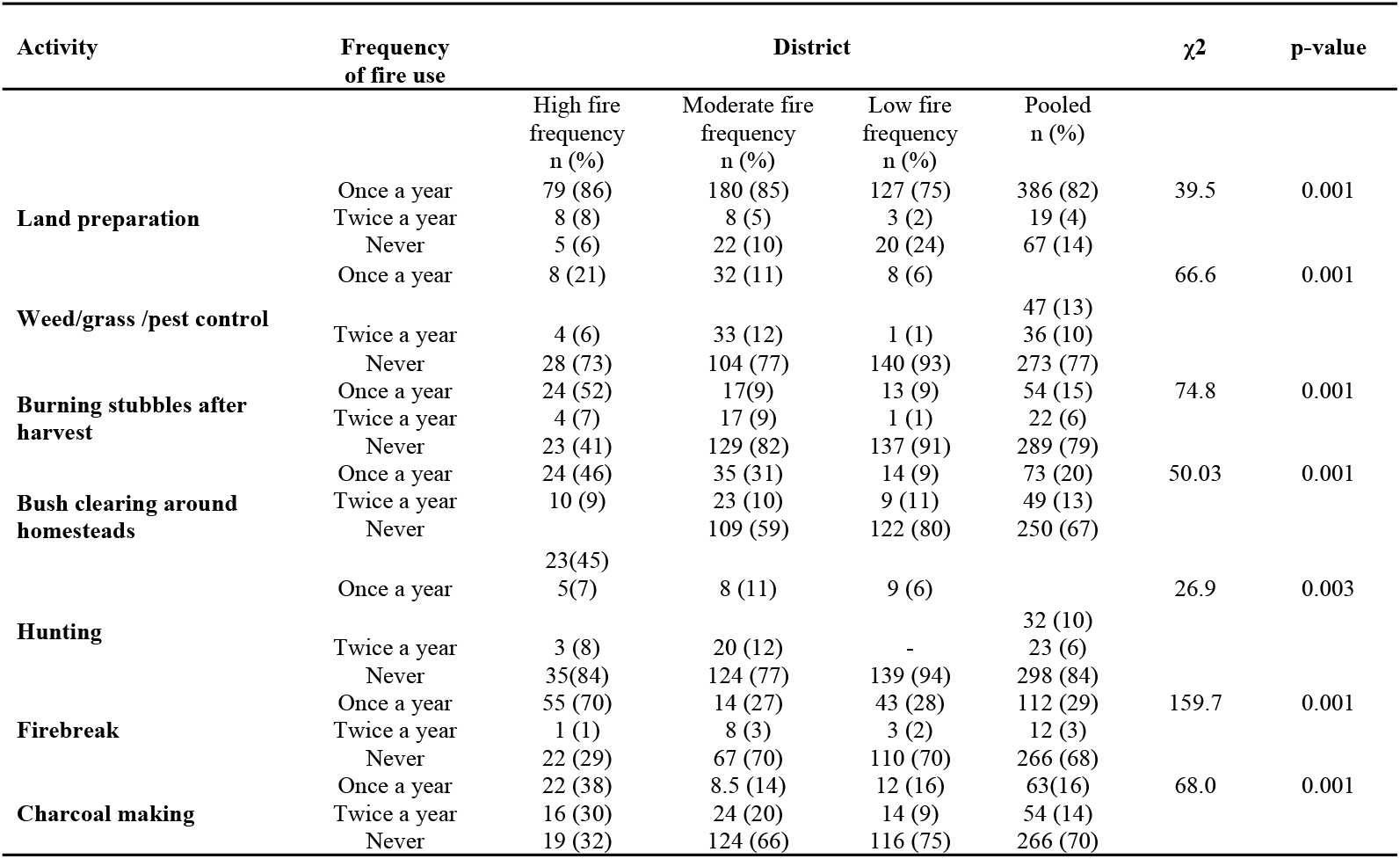
Respondents’ frequency of fire use for activities across the selected districts - high frequency district (6213-15254 counts) moderate fire frequency (2804-6213 counts) and low fire frequency (487-2804 counts) (Pearson’s chisquare, P < 0.01).

The was a significant association between the frequency of fire used for weed control and the district (χ^2^ = 66.6, df = 9, p < 0.001). On average, 13% of the respondents used fire once a year for weed control, while the majority (77%) never used fire for this purpose (Table 2).About a fifth (21%) of the respondents in the high-frequency zone, 11% in the moderate zone and 6% in the low zone, used fire once a year for weed control. In contrast, 73%, 77% and 93% of respondents in the high, moderate and low fire frequency zones, respectively, never used fire for weed control.

The majority of respondents (79%) never used fire to burn stubble after harvest, while less than a fifth (15%) used fire once a year. A little over half of the respondents in the high (52%) and low fire districts indicated that they burn stubbles after harvest. There was a significant association of frequency of fire use for stubble burning and districts (χ^2^ = 74.8, df = 9, p < 0.001).

Bush clearing around homesteads also showed strong association (χ^2^ = 50.3 df = 9, p < 0.001) amongst the districts. However, 67% of the respondents never used fire for clearing around homesteads. Twenty percent and 13% of the respondents used fire once and twice a year respectively, across the districts. However, about half (46%) of respondents in high fire frequency districts and as low as 9% respondents in low fire frequency district said they used fire once a year. Whereas 45% in high, 60% in moderate and 80% in low fire frequency districts never used fire for the same activity (Table 2).

Respondents’ use of fire to create firebreaks showed a strong significant association (χ^2^ = 50.33 df = 9, p < 0.001) among the districts. On average, the majority (70%) of the respondents in the moderate and low fire districts never used fire for fire breaks. Nearly three-quarters (70%) of the respondents in high fire districts, a little over a fourth of respondents in moderate (27%) and low (28%) fire frequency districts used fire once a year for fire breaks.

Similarly, there was a significant association of fire use for hunting (χ^2^ = 26.88, df = 9, p < 0.003) across the study districts. On average, 84% of the respondents across the districts never used fire for hunting, with as low as 7% in the high, 11% in the moderate and 6% in the low fire frequency districts used fire once a year (Table 2).

### 3.3 Fire control practice for selected activities

The majority of interviewees responded in the affirmative when asked whether they always controlled fire for land preparation and burning around homesteads. In contrast, the majority of respondents indicated they never controlled fire for the other selected activities (Table 3). Respondents who controlled fire and those who never controlled fire enumerated reasons for which fire used for the selected activities should be controlled. Thus, 62% of 532 mentioned that they control fire to prevent destruction to property, 29% said it was to prevent destruction to vegetation and wildlife, 6% mentioned to maintain soil fertility, and the remaining 3% indicated to prevent killing or injury to humans.

**Table 3:** Respondents’ practice of fire control across the fire gradient districts - high frequency district (6213-15254 counts) moderate fire frequency (2804-6213 counts) and low fire frequency (487-2804 counts) (Pearson’s Chi-square, P < 0.001).

| Activity | Practice of fire control | District | | | | $\chi^2$ | p-value |
| --- | --- | --- | --- | --- | --- | --- | --- |
|  |  | High fire frequency<br>n (%) | Moderate fire frequency<br>n(%) | Low fire frequency<br>n (%) | Pooled<br>n(%) |  |  |
| Land preparation | Always | 75 (79) | 125 (74) | 121 (73) | 321 (72) | 61.4 | **0.001** |
|  | Sometimes | 12 (14) | 15 (12) | 10 (6) | 57 (13) |  |  |
|  | Never | 6 (7) | 30 (15) | 34 (21) | 70 (16) |  |  |
| Weed//pest control | Always | 12 (15) | 19 (9) | 8 (6) | 26 (8) | 48.5 | **0.001** |
|  | Sometimes | 8 (10) | 18 (8) | 1 (1) | 29 (9) |  |  |
|  | Never | 3 (2) 7 (6) | 108 (83) | 132 (93) | 274 (83) |  |  |
| Burning stubbles after harvest | Always | 17 (59) | 4 (3) | 9 (7) | 30 (9) | 126.7 | **0.001** |
|  | Sometimes | 6 (7) | 5 (3) | 4 (3) | 15 (5) |  |  |
|  | Never | 28 (34) | 128 (95) | 129 (91) | 285 (86) |  |  |
| Bush clearing around homesteads | Always | 29 (65) | 30 (18) | 28 (20) | 87 (25) | 78.6 | **0.001** |
|  | Sometimes | 6 (8) | 7 (35) | 1 (1) | 15 (5) |  |  |
|  | Never | 22 (28) | 105 (79) | 114 (79) | 241 (70) |  |  |
| Hunting | Always | 9 (20) | 5 (2) | 6 (5) | 20 (6) | 67.4 | **0.001** |
|  | Sometimes | 2 (5) | 10 (5) | 2 (1) | 14 (4) |  |  |
|  | Never | 32 (75) | 131 (93) | 130 (94) | 293 (90) |  |  |
| Firebreak | Always | 42 (65) | 5 (6) | 45 (31) | 92 (27) | 171.9 | **0.001** |
|  | Sometimes | 3 (4) | 1 (1) | 1 (1) | 6 (2) |  |  |
|  | Never | 25 (31) | 115 (92) | 104 (68) | 244 (71) |  |  |
| Charcoal making | Always | 34 (64) | 21 (16) | 33 (23) | 88 (24) | 102.7 | **0.001** |
|  | Sometimes | 4 (6) | 12 (5) | 1 (1) | 17 (5) |  |  |
|  | Never | 24 (29) | 119 (79) | 110 (76) | 254 (71) |  |  |

There were significant associations between respondents’ fire control practices and the selected activities across the three fire gradient districts (χ2 = 61.4, df = 9, p < 0.001). Around three-quarters (72%) of the respondents indicated that they controlled fire for land preparation. Seventy-nine percent of the respondents in high fire frequency districts, and 74% for moderate and low (73%) indicated that they always controlled fire for land preparation (Table 3). Less than a quarter across the district never controlled fire during land preparation.

Contrary to the control of fire for land preparation, 82% of the respondents in each of the districts never controlled fire for weed control, but only a fifth of respondents in the high (10%) moderate (9%), and low (6%) fire frequency districts always controlled fire (χ^2^ = 48.5, df = 9, p < 0.001).

However, 59% of the respondents in the high fire frequency districts and less than a tenth of the respondents in the moderate (3%) and low (6%) districts, always controlled fire for burning stubble. However, the highest number (90%) of respondents in both moderate and the lowes 34% in high fire frequency districts never controlled fire for burning stubble (χ^2^ = 39.5, df = 9, p < 0.001).

As much as twice the number of respondents (65%) in high fire districts always controlled fire created for firebreaks than those in low fire frequency districts (31%), with only 6% in the moderate fire frequency districts (χ^2^ = 159.7, df = 9, p < 0.001).

The highest number of respondents in all the districts indicated that they never controlled fire for hunting, whilst less than a fifth of the respondents in the districts, controlled fire always (Table 3). During charcoal burning, respondents in the high fire (64%), moderate fire (16%) and low fire districts (23%) always controlled fire, whereas 26%, 79% and 76% in the high, moderate and low fire frequency districts, respectively, never controlled fire (Table 3).

### 3.4 Perceptions of the importance of fire regimes

There were varied perceptions of fire regimes. There was evidence of significant relationships in the perceptions of fire regimes in the high, moderate and low fire frequency districts (Table 4). Half of the respondents (50%) in the high fire frequency districts, 33% in the moderate and 43% in the low, were of the opinion that the season of burning was very important. While less than a fifth in each of the districts said the season of burning was not important.

**Table 4:** Respondents’ perceptions of the importance of fire regimes across the six districts

| Fire attribute | Importance of fire attributes | District | | | | $\chi^2$ | p-value |
| --- | --- | --- | --- | --- | --- | --- | --- |
|  |  | High fire frequency<br>n (%) | Moderate fire frequency<br>n (%) | Low fire frequency<br>n (%) | Pooled<br>n (%) |  |  |
| Season of burning (timing; dry, season, early or late) | Very Important | 46 (50) | 49(33) | 90 (53) | 185 (43) | 55.6 | 0.001 |
|  | Important | 28 (32) | 95(51) | 49 (29) | 173 (40) |  |  |
|  | Not important | 20 (18) | 23 (15) | 30 (18) | 73 (17) |  |  |
| Severity of fire (what is burned, soil vegetation mortality, high, moderate, low) | Very Important | 21(21) | 28 (17) | 30 (18) | 79 (18) | 40.8 | 0.001 |
|  | Important | 37(45) | 68 (41) | 47 (28) | 152 (36) |  |  |
|  | Not important | 33 (34) | 75 (43) | 8 (53) | 196 (46) |  |  |
| Frequency of fire (fire return interval and number of ignitions) | Very Important | 14 (18) | 11 (5) | 24(15) | 49 (12) | 53.8 | 0.001 |
|  | Important | 28 (30) | 62 (33) | 27(17) | 117 (29) |  |  |
|  | Not important | 47 (52) | 79 (63) | 112 (69) | 238 (59) |  |  |
| Type of fire (size and pattern) | Very Important | 20 (23) | 16 (7) | 19 (12) | 55 (13) | 37.7 | 0.001 |
|  | Important | 20 (22) | 62 (29) | 38 (23) | 120 (29) |  |  |
|  | Not important | 48 (56) | 85(63) | 105 (65) | 240 (58) |  |  |

The majority of respondents perceived the severity of fire as important but not very important: 45%, 41% and 28% in high, moderate and low respectively, indicated that the severity of fire was important. However, more than half of the number of respondents (53%) in the low fire districts, 33% of the respondents in high and 44% in moderate districts indicated that the severity of fire was not important.

About 59% of the respondents perceived that the frequency of burn was unimportant. However, 30% in high, 33% moderate and 17% low fire frequency districts said the frequency of burning was important. Similar to the responses on perceptions of fire frequency, on average the majority of respondents (58%) in each of the districts were of the opinion that the type of fire was unimportant. Thus, less than a quarter of the number of respondents in each of the districts thought the type of fire was very important (Table 4).

### 3.5 Respondents’ perceptions of the effects of fire on the environment and the frequency of fire use

About 77% of the respondents were of the view that fire was not good for the environment, 8% thought fire use was good for the environment while the remaining 14% indicated they did not know. However, thirty-five percent of the respondents indicated that burning was good for the environment, said that burning increased soil fertility, followed by 28% who were of the view that it improved crop yield, while less than 1% said it was good for hunting. In contrast, 42% of respondents who said that burning was not good, were of the view that fire destroyed vegetation, while 33% were of the opinion that fire decreased soil fertility with as few as 1% stating it exposed animals to hunting (Figure 3).

**Fig. 3:**
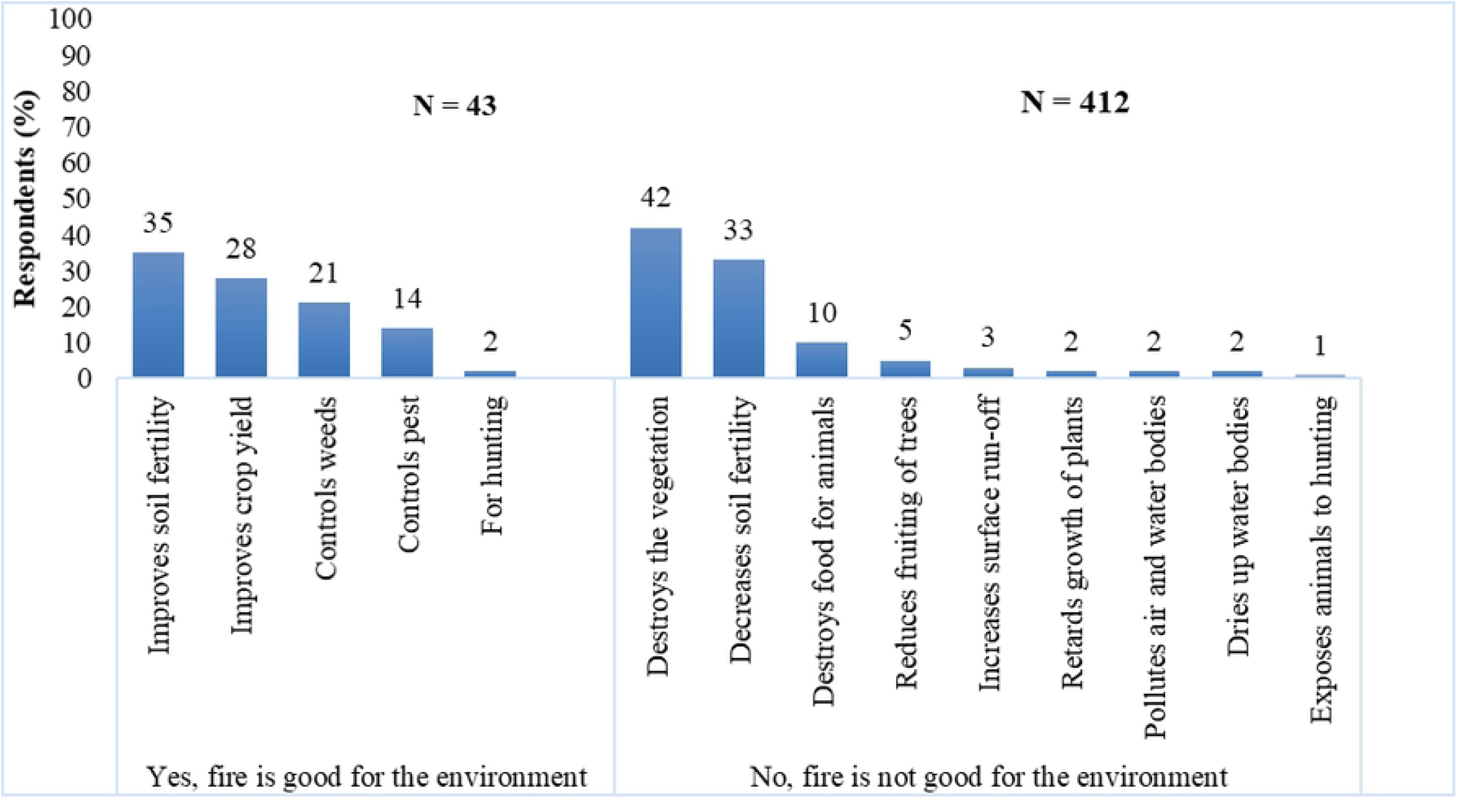
respondents’ opinion on the effects of fire on the environment

Generally, there were positive correlations amongst perception of fire effects on the environment and the seven cultural uses of fire. The strongest positive correlation was found between perceptions of fire effect on the environment and how often fire is used for land preparation (r = .140 n = 449, p = .003). However, a negative correlation was found between perception of fire effect on the environment and frequency of fire use for firebreaks, bush clearing around homesteads and hunting (Table 5).

**Table 5.** Respondents’ perceptions of fire effects on the environment correlated to how often fire is used for the seven socio-cultural practices.

|  |  | **In your opinion is fire good for the Environment?** | **How often do you use fire for land preparation for farming** | **How often do you use fire for hunting** | **How often do you use fire for weed control** | **How often do you use fire to clean farm after harvesting** | **How often do you use fire for bush clearing around homesteads** | **How often do you use fire for charcoal burning** | **How often do you use fire for fire breaks** |
| --- | --- | --- | --- | --- | --- | --- | --- | --- | --- |
| **In your opinion is fire good for the Environment?** | Pearson Correlation | 1 | .140** | .079 | .003 | .075 | -.015 | -.081 | -.005 |
|  | Sig. (2-tailed) |  | .003 | .152 | .954 | .161 | .773 | .129 | .929 |
|  | N | 501 | 449 | 333 | 336 | 347 | 354 | 356 | 375 |
| **How often do you use fire for land preparation for farming** | Pearson Correlation | .140** | 1 | .121* | .199** | .151** | .142** | .103* | .233** |
|  | Sig. (2-tailed) | .003 |  | .021 | .000 | .003 | .005 | .044 | .000 |
|  | N | 449 | 490 | 366 | 369 | 378 | 382 | 386 | 399 |
| **How often do you use fire for hunting** | Pearson Correlation | .079 | .121* | 1 | .146** | .238** | .288** | .209** | .154** |
|  | Sig. (2-tailed) | .152 | .021 |  | .005 | .000 | .000 | .000 | .004 |
|  | N | 333 | 366 | 370 | 362 | 359 | 358 | 356 | 350 |
| **How often do you use fire for weed /pest control** | Pearson Correlation | .003 | .199** | .146** | 1 | .126* | .043 | .113* | -.004 |
|  | Sig. (2-tailed) | .954 | .000 | .005 |  | .016 | .415 | .032 | .942 |
|  | N | 336 | 369 | 362 | 373 | 364 | 363 | 361 | 352 |
| **How often do you use fire to clean farm after harvesting** | Pearson Correlation | .075 | .151** | .238** | .126* | 1 | .494** | .256** | .247** |
|  | Sig. (2-tailed) | .161 | .003 | .000 | .016 |  | .000 | .000 | .000 |
|  | N | 347 | 378 | 359 | 364 | 382 | 370 | 367 | 360 |
| **How often do you use fire for bush clearing around homesteads** | Pearson Correlation | -.015 | .142** | .288** | .043 | .494** | 1 | .254** | .211** |
|  | Sig. (2-tailed) | .773 | .005 | .000 | .415 | .000 |  | .000 | .000 |
|  | N | 354 | 382 | 358 | 363 | 370 | 389 | 370 | 363 |
| **How often do you use fire for charcoal burning** | Pearson Correlation | -.081 | .103* | .209** | .113* | .256** | .254** | 1 | .366** |
|  | Sig. (2-tailed) | .129 | .044 | .000 | .032 | .000 | .000 |  | .000 |
|  | N | 356 | 386 | 356 | 361 | 367 | 370 | 392 | 363 |
| **How often do you use fire for fire breaks** | Pearson Correlation | -.005 | .233** | .154** | -.004 | .247** | .211** | .366** | 1 |
|  | Sig. (2-tailed) | .929 | .000 | .004 | .942 | .000 | .000 | .000 |  |
|  | N | 375 | 399 | 350 | 352 | 360 | 363 | 363 | 405 |

\*\* Correlation is significant at the 0.01 level (2-tailed). \* Correlation is significant at the 0.05 level (2-tailed)

## 4 DISCUSSION

### 4.2 Frequency of fire use for selected activities across the districts

The frequency of fire use is an important attribute of fire regimes which is characterised by the fire return interval and the number of fires that occur within a period. Most of the respondents were involved in farming as their occupation, hence, the high response rate for using fire once a year for land preparation. The response for fire use once a year corresponds with the unimodal rainy season and the long dry season (up to six months) in the study area. Studies have shown that crop farmers in the tropical savannas, burn to reduce thrash and get rid of the grass and other herbaceous vegetation from the previous rainy season to allow for hoeing or ploughing during land preparation for the next cropping (Amissah, Kyereh and Agyeman 2010; Shaffer 2010). The respondents who indicated that they used fire twice a year for land preparation, may do this on occasions where they have to clear a new place for cropping; the first burning is to clear the bush, then fire is used again to remove stumps of the large woody species (Eastmond and Faust 2006), as observed during the field reconnaissance survey for this study.

The high fire frequency districts are located in a closed savanna (Bagamsah 2005) which is characterised by vast areas of wildland of tall grasses and relatively good soil for agricultural production.. As a result, even people in the low fire areas, particularly those that are ‘urbanised’ move to these areas for farming using traditional agricultural practices, which could also contribute to high fire frequency. These districts are also characterised by high grass growth during the rainy season which enhances fuel load in the dry season. This results in high intensity fires with temperatures between 791°C and 868°C in the early season, and 790°C and 896°C in late season fires as observed by Kugbe, Fosu, and Vlek (2014) in the study area. Veenendal *et al.* (2018) also observed that fire return intervals are largest for tall savanna woodlands and dry forests. Thus, a possibility that fire in these areas may increase as conditions for fire ignition become more conducive for burning: heavy rainfall that promotes good grass growth and the hash Harmattan conditions (Amoako *et al.* 2018). Population growth in urban areas and the resultant high demand for pulses, cereals and yam grown in the agroforestry parklands (where smallholder farmers still use traditional methods of farming involving the use of fire) could also increase the use of fire for land preparation in West Africa.

According to Nyongesa (2018), the use of fire for weed control is another cultural practice amongst rural dwellers in most tropical savannas. Some respondents indicated that they used fire for weed control including the control of *Striga hermonthica.* This was also observed in a study in Kenya, where farmers indicated burning and hand-pulling to control *Striga hermonthica* on crop fields (Atera, Itoh, Azuma and Ishii 2018). Other studies (DiTomaso and Johnson 2006; Morton *et al*. 2010; Mckell, Wilson and Kay 2015), have also reported that the use of prescribed fires (also referred to as flame weeding) to control recalcitrant pests, weeds and invasive species has been successful with species such as *Centauries solstitialis* (Yellow star-thistle) in California (DiTomaso and Johnson 2006). Prescribed fire use in Ngorongoro, Kenya was also successful in the control of ticks (Fyumagwa *et al.* 2007). However, the use of weedicides and herbicides is probably gaining ground because of their ready availability in local markets and in remote areas of Ghana (Nkansah 2014). Some respondents explained that it is difficult to apply fire while crops are still in the field and would prefer to use chemicals and rather burn for land preparation. This could also be the reason for the relatively high responses for fire use after harvesting for burning stubble, which sometimes forms part of the process of land preparation and crop production (Chan and Heenan 2005).

Studies (Shaffer 2010; Nyongesa and Vacik 2018) have shown that the use of fire for clearing bush around homesteads during the dry season is a common practice in most rural communities in Africa. Most of the study districts are rural with homes having bushy surroundings which dry up every Harmattan season; as a result, applying fire is much cheaper and more practical than weeding and/or applying weedicides, as explained by some of the respondents. In contrast, the districts in urban areas with high human populations have more paved surroundings, thus little use of fire for this activity has been observed (Archibald 2016). Nyongesa and Vacik (2018) also have shown that the use of fire in crop fields is to save on labour and the cost of chemicals.

Homesteads are mostly used for compound gardens in the rainy season which supply vegetables to supplement the nutritional needs of the household. These supplementary farms are known as compound farms which are burnt to get rid of the dried stubble in time for the next rainy season. Burning around homesteads is also done to get rid of ticks and reptiles as these homesteads contain stables for cattle and other livestock (Mistry *et al.* 2018).

The practice of using fire for creating firebreaks was relatively high across the study area, which confirms the findings of Shaffer (2010) and Nyongesa and Vacik (2018), who observed in Mozambique and Kenya respectively, that using fire for firebreaks was done to reduce fire risks within the communities and to protect livestock as well as crops which are yet to be harvested.

Hunting and charcoal burning were sensitive items in the questionnaires, particularly in communities with government protected forests with more burned areas, either due to uncontrolled fires from hunting or poaching, as well as charcoal production (Walters 2012). The ‘burn to flush out’ method of hunting used by most people in the dry season is difficult to control and not recommended for hunting (MLFM 2006). Hence people would not indicate that they were involved in hunting. Generally, there are more farmers than hunters in these communities except for communal hunting where more (60 - 100) people are involved (Adongo et al. 2012; Abukari and Mwalyosi 2018)

In spite of the low response rate for questions regarding charcoal burning and hunting across the districts, anecdotal evidence, satelite observations (SERVIR West Africa 2019) and field observations during the data collection period revealed that there is a high usage of fire for charcoal production and hunting. We also observed a lot of charcoal production going on in some of the communities. The respondents’ reluctance in giving honest answers on their use of fire for charcoal production and hunting could be attributed to people’s awareness of the dangers and adverse effects that these activities have on the environment, human lives and property. Also, there has been a campaign against these practices on the radio, as well as local announcements made by traditional authorities at the beginning of every dry season (Amoako *et al.* 2015).

In addition, the respondents seem to be aware of the suggested policy to ban charcoal production and so most of the respondents would not indicate that they used fire for charcoal burning. It is not surprising that people’s perception and the frequency of fire use for this activity showed a negative correlation. We realised that although people were aware of the effects of fire on the environment, they still produce charcoal, which has become an alternative source of livelihood income in the dry season in recent times due to the unpredictable rainfall and crop failure. Agyeman *et al.* (2012) suggested that instead of placing a ban on charcoal which is a lucrative source of livelihood income for rural people in the savannas of Ghana, the government should put in place measures to sustain the charcoal industry. However, Msuya, Masanja and Temu (2011) argued that that efforts should be made to reduce and prohibit the use of charcoal in cities and towns in order to reduce the degradation and its attendant effects on the savanna and forests in Africa.

#### 4.2 Fire control practice

We defined the practice of fire control for the selected activities as the use of fire with caution and supervision to achieve the desired results. Shaffer (2010) indicated that fires that are deliberately ignited for livelihood activities are usually controlled. Bowman, Amacher and Merry (2008), also found that traditional households that use fire in Brazil in the Amazon were actively engaged in fire management to protect their properties and farmlands.

It was observed that the high fire frequency districts had the highest responses for the use of fire once a year for almost all the selected activities. The respondents also exhibited more knowledge and practice of fire control for all the activities than those in moderate and low fire frequency districts who never controlled fire. However, this may not necessarily mean that those who never control fires are not aware of the control measures, but may use less fire hence; there is little need for fire control. The high proportion of respondents who control fire for the selected activities in the high fire frequency districts is an indication that the people were aware of the need to control fires to prevent destruction to humans, wildlife and property. This has also observed in other studies in Brazil (Bowman, Amacher and Merry 2008) and Kenya (Nyongesa and Vacik 2018) where fires that are applied intentionally are always controlled.

#### 4.3 Perceptions of the importance of fire regimes and the knowledge of fire effects on the environment

It was noted that fire regimes in the Guinea savanna are characterised by land use patterns and practices linked to livelihood and socio-cultural activities, and which are in turn determined by the season of burning, severity, frequency and size. The season of burning was rated very important, which confirms why most burning is done in the dry season to prepare farm lands for cropping in the rainy season (Veenendaal et al. 2018). This finding agrees with other studies which found vegetation burning in tropical savanna (Mistry, 1998) to mostly occurs in the dry season; the active fire season in the Sudano Guinean savanna (Rose Innes 1972; Dwomoh and Wimberly 2017; Laris *et al.* 2017). The same studies (Rose Innes 1972; Dwomoh and Wimberly 2017; Laris *et al.* 2017) reported that most of the fires are caused by humans for the activities mentioned in this study and other studies (Archibald 2016; Dwomoh and Wimberly 2017). Another important attribute which was explored was the time of burning within the dry season. Our observation during a reconnaissance survey was that most burning was done during December and January (early dry season) with a reduction in the number of fire occurrences towards the mid (February) and (March-April) late dry season. Respondents’ explanation indicated that burning is, however, dependent on the time the rains end (drought sets in). Thus, the timing and severity of drought within the dry season are fire conditions that play a major role in the fire regimes of the Guinea savanna.

Most respondents indicated that fire could be devastating to all aspects of the environment, including pollution of water bodies, soil erosion if not controlled; however, they attached more importance to the season of burning over the severity/intensity of the fire. For instance, Laris *et al*. 2017, also placed more emphasis on the season of burning than severity. The authors attributed the season of burning as a determinant of intensity and severity. Nonetheless these attributes are characterised by the extent of drought which influences fuel load, moisture content and local conditions (land use patterns and practices) and vary from place to place in the Guinean savanna of West Africa. Gambiza *et al*. (2005) and Govender, Trollope and van Wilgen (2006) also observed that in Southern Africa that fire severity is determined by the season of burning which is also influenced by factors including the moisture content, fuel load and fuel characteristics. A recent study by N’Dri *et al.* (2018) on fire behaviour in Guinea savanna, confirmed that the severity and intensity of fire is influenced by season of burning.

However, most of the respondents did not see the vital link between the season of burning and the severity of fire. Although the respondents indicated that they burned earlier in the dry season when the vegetation is not too dry, they rated severity of fire as not important. This suggests a knowledge gap in respondents understanding of the concept of fire regime. This agrees with Rodríguez-Trejo *et al*.’s (2011) commendation that that results of scientific research on the concept of fire regimes should be merged with traditional fire knowledge for long term integrated fire management.

Fire frequency was rated highly by respondents as unimportant. There was however a strong positive correlation between fire effect on environment how often people used fire for land preparation. Contrarily, the majority of respondents did not perceive fire return intervals and number of ignitions as a problem. Minnich *et al*. (1993) claimed that human-caused fire frequency (number of ignitions) is unimportant, because the onset of natural fires could be more devastating since natural fires occur when there is so much fuel load for burning, resulting in high intensity fires than frequent human ignited fires. However, Griffiths, Stephen and Garnett (2015) found that fire frequency has an enormous effect on both plant and animal species, and therefore should not be underestimated.

Most respondents thought the type of fire which encompasses pattern and size of fire was unimportant. However, Govender, Bond and van Wilgen (2006), have shown that fire severity is influenced by the type of fire which is in turn affected by the season of fire.In contrast, Govender et al (2006) have shown that fire severity is influenced by the type of fire which is in turn affected by the season of fire. This suggests the need to address the knowledge gap in managing the cultural uses of fire and fire regimes in savanna.

As shown in our study, most fires in these savannas occur annually and at specific times, thus have defined the anthropogenic fire regimes in these savannas (Archibald 2016). There is a need, for further studies to understand the complexities of human-driven fire regimes in Africa savannas.

## 5 CONCLUSION

The study confirms that the use of fire for agriculture and some socio-cultural purposes is a characteristic of traditional livelihood practices, which are mostly subsistence, but could have enormous impacts on the environment, as outlined by the respondents. Most of the respondents are farmers who use fire once a year for land preparation for cropping. The high fire frequency districts used fire for almost all the selected activities, thereby increasing the fire occurrences in these districts, which confirms the fire count data obtained from the CSIR Meraka Institute.

The study revealed that fires that are set on purpose were mostly controlled, particularly for land preparation for cropping, which shows that people are aware of the hazards associated with uncontrolled fires. There’s the need to study the ecological effects of traditional fire use.

The season of burning was highly rated as a very important component of fire regimes. The respondents’ stated that early and late season burning was dependent on the time the drought sets in. Notwithstanding, the type of fire was the least rated amongst the four attributes of a fire regime. More education on the impacts of different fire regimes could improve fire management in fire-prone areas. Most respondents were aware of the adverse on the effects of fire on the environment. This positive knowledge base could also be a foundation for the education and training in good fire management practices in the rural communities of West Africa.

The fire volunteer squads in communities mentioned in Amanor’s (2002) paper on bushfire management, culture and ecological modernisation in Ghana should be retrained and revived by district assemblies to assist in the management of fires for agricultural purposes as well as other socio-cultural practices within rural areas in the north of Ghana where fire use is rampant. Nonetheless, traditional authorities have the mandate of influencing cultural and traditional knowledge systems and must be recognised in collaborations among stakeholders to improve perceptions and fire use practices. Accordingly, collaborative sharing and learning of traditional ecological fire knowledge management between regional, district directorates of Ministry of Food and Agriculture, the Forestry Commission, the Environmental Protection Authority, Environmental NGO’s and researchers is important in the management of the savannas of the Guinea savanna and West Africa at large.

## Acknowledgements

Our profound gratitude goes to the leaders of the study community for their support. We are very grateful to the research assistants of the Faculty of Natural Resources and Environment, University for Development Studies, Tamale, Ghana, who assisted in the collection of field data.

Our sincere appreciation is extended to Dr Amos Karbo-Bah of the University of Energy and Natural Resources, Sunyani, Ghana, who provided the first batch of data on fire counts; as well as to Mr Ndumiso Booi of the CSIR Meraka Institute, South Africa, who provided a five-year data set on daily fire counts for Ghana. The financial support from the Organisation of Women in Science in the Developing World (OWSD) during my PhD Fellowship is very much appreciated.

## Author Contributions

Esther Ekua Amoako and Professor James Gambiza designed this research: Esther carried out the field research, analysed the data and wrote the manuscript. Professor Gambiza made inputs by comments, read and approved the final manuscript.

